# Homeolog expression quantification methods for allopolyploids

**DOI:** 10.1101/426437

**Authors:** Tony Kuo, Masaomi Hatakeyama, Toshiaki Tameshige, Kentaro K. Shimizu, Jun Sese

**Affiliations:** Artificial Intelligence Research Center, AIST, 2-3-26 Aomi, Koto-ku, 135-0064, Tokyo, Japan; AIST-Tokyo Tech RWBC-OIL, 2-12-1 Okayama, Meguro-ku, 152-8550, Tokyo, Japan; Department of Evolutionary Biology and Environmental Studies, University of Zurich, Winterthurerstrasse 190, CH-8057, Zurich, Switzerland; Functional Genomics Center Zurich, Winterthurerstrasse 190, CH-8057, Zurich, Switzerland; Swiss Institute of Bioinformatics, Quartier Sorge - Batiment Genopode, 1015 Lausanne, Switzerland; Kihara Institute for Biological Research, Yokohama City University, 641-12, Maioka, Totsuka-ku, 244-0813, Yokohama, Japan

## Abstract

Genome duplication with hybridization, or allopolyploidization, occurs in animals, fungi, and plants, and is especially common in crop plants. There is increasing interest in the study of allopolyploids due to advances in polyploid genome assembly, however the high level of sequence similarity in duplicated gene copies (homeologs) pose many challenges. Here we compared standard RNA-seq expression quantification approaches used currently for diploid species against subgenome-classification approaches which maps reads to each subgenome separately. We examined mapping error using our previous and new RNA-seq data in which a subgenome is experimentally added (synthetic allotetraploid *Arabidopsis kamchatica*) or reduced (allohexaploid wheat *Triticum aestivum* versus extracted allotetraploid) as ground truth. The error rates in the two species were very similar. The standard approaches showed higher error rates (> 10% using pseudo-alignment with Kallisto) while subgenome-classification approaches showed much lower error rates (< 1% using EAGLE-RC, < 2% using HomeoRoq). Although downstream analysis may partly mitigate mapping errors, the difference in methods was substantial in hexaploid wheat, where Kallisto appeared to have systematic differences relative to other methods. Only approximately half of the differentially expressed homeologs detected using Kallisto overlapped with those by any other method. In general, disagreement in low expression genes was responsible for most of the discordance between methods, which is consistent with known biases in Kallisto. We also observed that there exist uncertainties in genome sequences and annotation which can affect each method differently. Overall, subgenome-classification approaches tend to perform better than standard approaches with EAGLE-RC having the highest precision.

## 1 Introduction

Genome duplication, termed polyploidization, is widespread in plants with up to 35% of land plants being recent polyploids [1]. Many crop species in particular are allopolyploids [2], which involves the hybridization of two different species with genome duplication. Thus, there is much interest in the study of genome duplication and the advantages or disadvantages this phenomenon may convey. In order to explore the underlying mechanisms that may provide adaptation, many gene expression studies have been conducted on both natural and synthetic allopolyploid species [3, 4, 5, 6, 7, 8]. Allopolyploid species have traditionally been difficult to analyze at the whole genome scale due to the large size of their genomes and the high levels of sequence similarity between duplicated chromosomes. These duplicated gene copies, called homeologs, are in general highly similar and pose challenges to gene expression analyses. However, these homeologs and bias in their expression are of great interest because they potentially contribute to adaptation in polyploid species [9, 10, 11].

Recent improvements in long sequencing read technologies and linkage strategies [12, 13, 14] have allowed for breakthroughs in polyploid genome assembly. In plant biology especially, recent allopolyploid species such as bread wheat [15] as well as many other agriculturally important plant species have benefited [16, 17]. Now that *de novo* genome assembly for allopolyploids is no longer as formidable as it once was, a large number of polyploid reference genomes are expected to become available in the near future to facilitate genome wide studies. Accordingly, there is a need to evaluate expression quantification methods given the presence of homeologs in allopolyploids. It is possible that the high level of sequence similarity between homeologs may pose challenges to read mapping and consequently, expression quantification as well as other types of sequence analysis. Currently, it is unclear whether the expression quantification methods in use currently for diploids are suitable for allopolyploids due to the lack of studies examining this issue.

In this study we pose the question “can diploid RNA-seq methods be directly applied to allopolyploids?” and evaluate different approaches and methods for homeolog expression quantification in allopolyploids. To evaluate methods in polyploids, analysis on genetic materials with and without subgenomes is highly valuable as a form of ground truth. Here we used synthetic allotetraploid *Arabidopsis kamchatica* (Fisch ex DC.) K. Shimizu & Kudoh and performed tests with its two direct parental accessions of *Arabidopsis halleri* and *Arabidopsis lyrata*. For hexaploid wheat *Triticum aestivum* Chinese Spring [18] with *ABD* subgenomes, we performed tests with tetra-Chinese Spring (AB subgenomes) where the *D* subgenome was experimentally removed [19] as well as *Aegilops tauschii* the diploid progenitor of the wheat *D* subgenome.

We test, to the best of our knowledge, all known approaches to quantify expression in polyploids with four approaches:

1. A standard genome alignment based RNA-seq analysis on the full allopolyploid reference genome with two different alignment tools STAR [20] and LAST [21, 22].
2. A pseudo-alignment based method with Kallisto [23] on the full allopolyploid transcriptome.
3. A subgenome-classification approach with HomeoRoq [7], which maps read sequences to each subgenome separately.
4. A subgenome-classification approach with EAGLE-RC [24], which maps read sequences to each subgenome separately and also explicitly uses genotype variations that discriminate between homeologs as constraints in analysis.

Our results show that EAGLE-RC had the lowest error rate (*A. kamchatica*: 0.40%, hexaploid wheat: 0.49%) for alignments to the correct subgenome, while Kallisto had the highest error rate (*A. kamchatica*: 12.43%, hexaploid wheat: 13.44%). LAST and STAR had similar error rates in *A. kamchatica* but LAST was more precise in hexaploid wheat. In general, performance between methods were comparable for *A. kamchatica* but not for hexaploid wheat. We also observed systematic differences in low expression genes that impacted the homeolog expression bias results. Other concerns include uncertainty in the completeness and accuracy of the genome sequence and annotation, which affected each method differently due to differences in constraints. This may be especially relevant as polyploid species have only begun to be sequenced and assembled in large numbers and the gene annotations are in their first iterations. In the face of this uncertainty, EAGLE-RC is the most precise in our evaluations.

## 2 Methods

### 2.1 RNA sequence data and reference genomes

We evaluated methods on two allopolyploid species, tetraploid *A. kamchatica* and hexaploid wheat *T. aestivum*.

The natural species *A. kamchatica* [25, 26] was derived from two diploid species *A. halleri* and *A. lyrata* recently [27, 8]. It is a model polyploid with a broad distribution range, self-compatibility and transformation technique [27, 28]. To construct synthetic polyploids, we used two highly homozygous parental accessions used for genome assembly: *A. halleri* Tada mine W302 (ver 2.2, scaffolds N50 712 kb) [29] and *A. lyrata* lyrpet4 (ver 2.2, scaffolds N50 1.2 Mb) [8]. The two genotypes were crossed then the genome doubling was induced by colchicine treatment. Although synthetic polyploidization may occasionally activate transposable elements or induce chromosomal rearrangements, the subgenomes of the synthetic polyploid were derived from the merging of the two parental genomes and are highly or completely identical, providing a unique opportunity to evaluate RNA-seq methods in allopolyploids.

In order to assess classification accuracy in synthetic *A. kamchatica*, we used data from the parental species so that the ground truth of a read’s subgenome origin is known. *A. halleri* subsp. *gemmifera* and *A. lyrata* subsp. *patraea* RNA-seq data [30] was obtained under DDBJ accession DRP003263 submission DRA004364. Briefly, this dataset consists of four samples each for *A. halleri* and *A. lyrata* with 2 × 100 bp paired-end reads for a total of approximately 20 Gb and 21 Gb respectively. For differential homeolog expression analysis, synthetic allotetraploid *A. kamchatica* RNA-seq data [7] was obtained under DDBJ accession DRP01140. Briefly, this dataset consists of three biological replicates of *A. kamchatica* before and after cold stress with 2 × 100 bp paired-end reads for a total of approximately 12 Gb and 10 Gb respectively.

The whole genome assembly and annotation for hexaploid wheat *T. aestivum* with *ABD* subgenomes was obtained from the International Wheat Genome Sequencing Consortium [18] (assembly ver 1.0 and annotation ver 1.0, N50 22.8 Mb). The assembly quality is at the chromosome level and the reference genome was split into *A*, *B* and *D* subgenomes allowing for separate read mapping in subgenome-classification methods.

In order to assess classification accuracy in wheat, we utilized tetra-Chinese Spring in which the *AB* subgenomes were extracted by removing the *D* subgenome by repeated backcrossing [19]. Thus, the genome sequence of tetra-Chinese Spring must be very close to the *AB* subgenomes of hexaploid Chinese Spring.

Because there is no genetic material of extracted *D* genomes from hexaploid wheat, we used *A. tauschii* KU-2076 (resource in Kyoto University, collected in Iran). The *D* subgenome of hexaploid wheat is known to be derived from this species, thus it should be highly similar though there is divergence due to within-species variations [31]. We obtained RNA-seq data of tetra-Chinese Spring and *A. tauschii* in triplicate for a total of 2.1 Gb and 2.8 Gb respectively [data submission in progress]. For differential homeolog expression analysis, *T. aestivum* RNA-seq data was obtained from NCBI (BioProject PRJEB12358) with SRA accessions ERR120175[2-4] and ERR120177[0-2] describing samples, in triplicate, 24 hours after inoculation of fungal pathogen *Fusarium graminearum* and mock inoculation for a total of 17.3 Gb and 16.1 GB, respectively.

### 2.2 Plant growth, RNA isolation, and sequencing

The plants were grown at 16°C in 8h light / 16h dark cycle with 60% relative humidity for two weeks and leaf tissues were harvested. RNA was extracted from each tissue using RNeasy Plant Mini Kits (QIAGEN, Hilden, Germany) in combination with DNase I treatment (QIAGEN). Illumina sequencing libraries were made by TruSeq Stranded mRNA Library Prep Kit. RNA-seq was conducted using Illumina HiSeq 4000 at the Functional Genomics Centre, Zurich.

### 2.3 Homeolog identification

To annotate homeologs in *A. kamchatica*, we constructed RNA transcripts from the gene models in the *A. halleri* and *A. lyrata* gene annotations. Homeologs were then identified based on reciprocal best hit for each subgenome’s transcripts. We required hits to have E-value less than 10^−10^ with at least 200 aligned bases in both transcripts, resulting in 24, 329 homeolog pairs identified.

To annotate homeologs in *T. aestivum*, we constructed RNA transcripts from high confidence gene models belonging to the *A*, *B*, and *D* subgenomes, including the UTR regions. We then identified pair-wise homeologs through reciprocal best hit for combinations *AB*, *AD*, *BD* and then triple copy homeologs by checking for genes in *AB* that share the same hit for *D* in their respective *AD* and *BD* hits, resulting in 21,196 triple copy homeologs identified.

### 2.4 Standard RNA-seq analysis

We tested a standard genome alignment based RNA-seq expression quantification approach (Figure 1a) aimed at differential expression analysis [32] that is often used for diploids:

**Figure 1:**
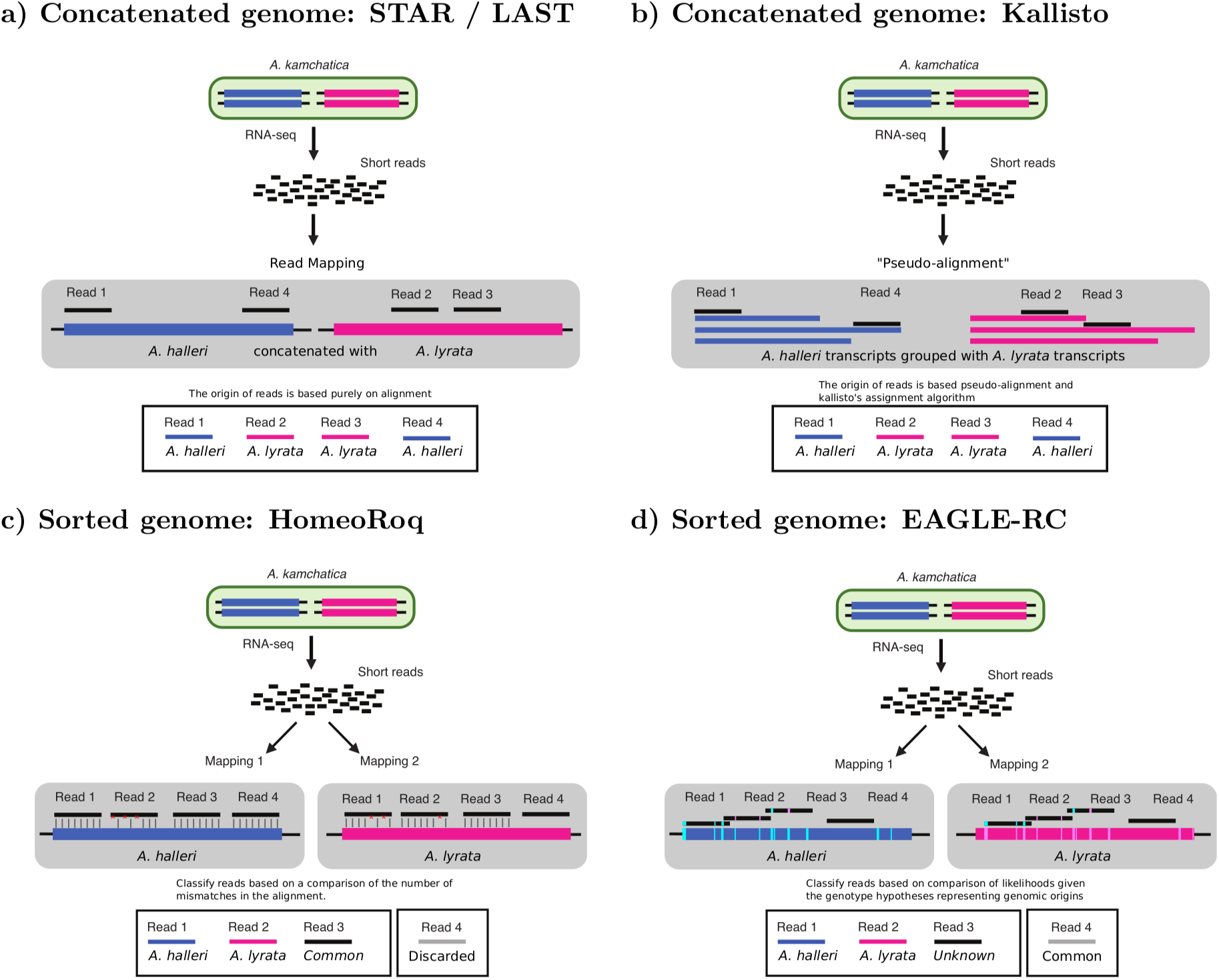
We compare different four approaches for quantifying homeolog expression: **a)** a standard genome alignment based RNA-seq analysis on an allopolyploid (concatenated) reference genome using alignment tools STAR and LAST. **b)** a pseudo-alignment workflow using Kallisto that is performed on the (concatenated) transcripts of the allopolyploid. **c)** a subgenome-classification analysis that performs alignment on each subgenome’s reference separately, then performs read classification based on number of mismatches using HomeoRoq. **d)** a subgenome-classification analysis that performs read alignment on each subgenome’s reference separately, then performs read classification based on the likelihood of the read to the genotype using EAGLE-RC, discarding common reads.

1. Map reads to the allopolyploid reference genome (STAR (ver 2.5.2b) [20], LAST (ver 809) [21, 22]).
2. Count reads using featureCounts [33] at the transcript level.
3. Extract homeolog specific read counts.

To construct the reference genome of *A. kamchatica*, we concatenated the reference genomes of its two parental species, *A. halleri* and *A. lyrata*, to obtain an allopolyploid reference. For LAST, we also filtered out read alignments with MAPQ scores less than 20, while STAR MAPQ scores are not as useful for thresholding. In *T. aestivum*, we excluded all reads that mapped to chrUn.

We tested a pseudo-alignment RNA expression quantification workflow (Figure 1b) using Kallisto [23], which is also often used for diploids. The built-in expression quantification in Kallisto was used to count reads (*est*_*counts*) at the transcript level. When evaluating classification accuracy, we used the pseudobam option in Kallisto to output read assignments and evaluate the proportion which were misassigned.

### 2.5 Subgenome-classification analysis

We tested a subgenome-classification approach where, in contrast to the standard approach, the sequencing reads are mapped separately to each subgenome of an allopolyploid’s reference genome (Figures 1c, 1d) using STAR. Then we utilized a read classifier (HomeoRoq, EAGLE-RC) to assign reads to their subgenome origin, if possible.

HomeoRoq [7] (Figure 1c) classifies reads based on the number of mismatches, up to a maximum of 10, between the read and the genome sequence of each subgenome, where reads must be mappable to both subgenomes in order to be considered. In contrast with EAGLE-RC, which requires computing the subgenome-discriminating variants, HomeoRoq does not require comparative analysis between different subgenomes in advance of classification.

The basic EAGLE model [24] is a generative model for read sequences used to calculate the likelihood of a read given a reference genotype hypothesis and an alternative genotype hypothesis. In EAGLE-RC (Figure 1d), the basic model was extended to perform read classification using the variants that discriminate between homeologs. During homeolog identification, subgenome specific genotype differences (i.e. variants) that discriminate between homeologs are determined. For *A. kamchatica*, reads are mapped to the reference genomes of *A. halleri*, *H*, and *A. lyrata*, *L*, separately. EAGLE-RC then calculates the probability given each subgenome as the reference hypothesis *G*_ref_ and the other subgenome as the alternative hypothesis *G*_alt_:

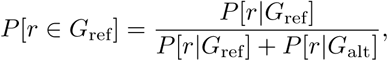

where the classification is determined by the reference with the highest probability, requiring the winning hypothesis to be at least probability 0.95 with marginal probability at least 0.51, otherwise it is “unknown”, where the marginal probability is the proportion of the winning hypothesis over the sum of all subgenome as the reference hypotheses.

For hexaploid *T. aestivum* subgenome-classification, we performed a bottom-up workflow using a series of pair-wise classifications (Figure 2). For HomeoRoq, we determined, if possible, the consensus classification from pair-wise classifications. For example, a read is classified as *A* if there is a consensus *A* classification in both *AB* and *AD* pair-wise classifications comparisons.

**Figure 2:**
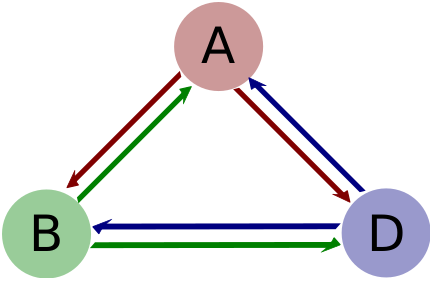
We perform read classification on hexaploid *T. aestivum* using a bottom-up approach from a series of pair-wise classifications with the *A*, *B*, and *D* subgenomes. A final classification from pair-wise analysis is obtained via consensus for HomeoRoq and via highest probability for EAGLE-RC.

For hexaploid subgenome-classification with EAGLE-RC, the pair-wise likelihoods per read were calculated, where for each reference hypothesis *G*_ref_, there are two alternative genome hypotheses *G*_alt1_ and *G*_alt2_.

The probability of a read belonging to a given reference is then:

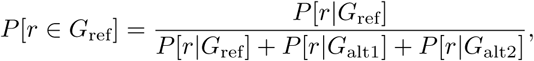

where the classification is determined by the reference with the highest probability, requiring the winning hypothesis to be at least probability 0.95 with marginal probability at least 0.51, otherwise it is “unknown”.

### 2.6 Differentially expressed homeologs

To identify differentially expressed homeologs (DEH) in *A. kamchatica*, we used DESeq2 [34] on the read count data for the *A. halleri*-derived subgenome and *A. lyrata*-derived subgenome separately. We required a homeolog to be differentially expressed with 0.05 or better False Discovery Rate (FDR) in at least one of the subgenomes. We then performed Fisher’s exact test on the combined read counts to determine significant differential homeolog expression ratio (p-value < 0.05 and fold change ≥ 2). This definition of differential expression requires significant change in gene expression in at least one subgenome along with the homeolog expression ratio under different conditions.

To find DEH in hexaploid *T. aestivum*, we used DESeq2 on the read count data for each *A*, *B*, and *D* subgenomes separately, requiring a gene to be differentially expressed with 0.05 or better FDR in at least one of the subgenomes. We then performed a series of Fisher’s exact test on the combined read counts (*A* vs *BD*, *B* vs *AD*, *D* vs *AB*) to test for differential homeolog expression ratio. A homeolog is differentially expressed if at least one test has p-value < 0.05/3 and fold change ≥ 2.

## 3 Results

### 3.1 *A. kamchatica* read classification

Generally, sequence analysis is based on how accurately read sequences can be mapped to the reference. In the case of allopolyploids, read alignment must deal with a higher degree of repetitiveness than in diploids due to homeologs.

We tested the standard genome alignment approach using the widely used RNA-seq read mapping tool, STAR. Though this tool is often used in diploid read mapping, it is unclear whether it will be suitable for allopolyploids. Unfortunately, the mapping quality score from STAR is not suitable for thresholding “uniqueness” due to how it assigns scores in an almost binary manner. Thus we also tested LAST, a general sequence alignment tool that can estimate the probability that an alignment represents the genomic source of the read. For example, a given read aligns to a single location with no mismatches, but aligns to five other locations with one mismatch each. This read may be deemed a unique best hit, but there may be a reasonable probability, depending on read length, that it came from any of the other alignments with a single base-calling error. LAST allows us to set a degree of uniqueness (i.e. 0.05 mismap probability) as a cut-off threshold that is convenient for handling this type of uncertainty.

We examined tetraploid *A. kamchatica* first because it may be less complicated than hexaploid wheat, which we describe later in this study. Here, we evaluated the accuracy of each tool and each approach by how well they assigned reads to the correct subgenome. For *A. kamchatica*, the ground truth is known by testing with pure *A. halleri* and *A. lyrata* RNA-seq data. Table 1 shows the classification performance of each approach for *A. kamchatica*. One point to note is that the mapping rate is affected by alignments to non-homeologs, thus we also showed the number of homeologs with detected expression in our evaluations.

**Table 1:** Classification performance for *A. kamchatica*. Results are averaged across 8 samples, 4 each for *A. halleri* and *A. lyrata*. The percent mapped refers to the number of reads that were mappable and in the case of subgenome-classification, the number of reads mappable to any subgenome. The classification error refers to the proportion of mapped reads which were assigned to the wrong subgenome. The number of expressed homeologs with > 1 read in any sample are also shown for each subgenome.

|  |  | Classification error |  | A. halleri | A. lyrata |
| --- | --- | --- | --- | --- | --- |
|  | Mapped | H to L | L to H | Homeologs | Homeologs |
| STAR | 93.40% | 11.09% | 11.40% | 21979 | 22030 |
| LAST | 71.59% | 2.87% | 2.50% | 21918 | 21990 |
| Kallisto | 92.22% | 12.42% | 12.44% | 23195 | 23234 |
| HomeoRoq | 93.67% | 1.64% | 1.23% | 21722 | 19653 |
| EAGLE-RC | 93.67% | 0.48% | 0.33% | 21385 | 21555 |

We calculated the precision by using classification error rate as the criteria. It is clear that subgenome-classification approaches (HomeoRoq and EAGLE-RC) performed better than the standard alignment based approaches (STAR, LAST, Kallisto) using a concatenated genome. In the standard alignment based approach using a concatenated reference genome, LAST’s mismap probability model was seen to be beneficial to read classification showing a much lower classification error rate than STAR.

Kallisto showed the lowest precision among all methods, though it showed the highest number of expressed homeologs detected. That Kallisto’s performance was the most affected by the presence of homeologs is perhaps due to a reduction in the number of unique kmers, relative to diploid analysis, which is essential for the method to find unique read to gene associations. To quantify the reduction in the number of unique kmers in a tetraploid reference relative to diploid reference, we performed a simulation with 1000 trials of 100 randomly selected *A. halleri* genes with randomly assigned SNPs at varying degrees of sequence divergence (Table S1). This analysis shows that there is a large reduction in the number of unique kmers given one extra gene copy depending on the pair-wise sequence divergence. For a point of reference, *A. kamchatica* is estimated to have approximately 2-3% divergence between homeologs [7].

EAGLE-RC showed the highest precision in read classification though it had a lower number of expressed homeologs detected. HomeoRoq was less precise than EAGLE-RC while having a similar number of expressed homeologs detected. The main difference in methodology between EAGLE-RC and HomeoRoq is that EAGLE-RC utilizes genotype information explicitly while HomeoRoq relies on comparing the number of mismatches, implicitly comparing genotype differences. However, simply comparing the number of mismatches is susceptible to spurious mappings, because a read alignment with 9 versus 10 mismatches favors the one with 9 mismatches to the same degree as it would favor an alignment with 0 versus 1 mismatch.

Another point to consider is that the quality of the reference may differ between subgenomes and there may be uncertainty from missing regions or erroneous annotations. HomeoRoq requires reads to be mappable to both references, which constrains read counting to genome regions that exist in both subgenomes. However, this comes with the disadvantage that more divergent regions may be excluded due to reads being mappable to only one homeolog. EAGLE-RC require reads to be mapped to regions that are different between homeologs and thus can be classified based on those differences, which is constrained to regions in the homeologs’ gene models that can be pair-wise aligned in order to compute the genotype differences between homeologs. We performed a simulation analysis where reads were simulated using ART [35] from annotated gene models with no divergence from the reference and calculated the classification performance (Table S2). The results show that even in this ideal scenario, there were reads which could not be classified with certainty by LAST, HomeoRoq, and EAGLE-RC due to reads mapping equivalently to both subgenomes. It also shows that in ideal conditions, STAR and Kallisto, despite higher error rates, were excellent in their true positive rates. However, the ideal condition of data with no divergence from reference, no sequences outside of annotated genes, and perfectly reflect gene models is not realistic in practice.

### 3.2 *A. kamchatica* homeolog expression quantification

Though read mapping is the foundation, downstream read counting methods and differential expression analysis may potentially be able to correct for artifacts or ambiguity in read alignments. We revisited classification error using the quantified read counts for homeologs in *A. kamchatica* (Table 2), although we suggest some caution in the interpretation of the number of reads quantified and thus the classification error rate in this result as HomeoRoq and EAGLE-RC have additional expression quantification processes to count subgenome-common reads. For Kallisto, we used the estimated read count from its output while for all other methods, we obtained read counts using featureCounts.

**Table 2:** Error rate for *A. kamchatica* using quantified read counts, averaged across 3 samples each for *A. halleri* (H) and *A. lyrata* (L).

|  | Classification error |  | Reads quantified |
| --- | --- | --- | --- |
|  | H to L | L to H |  |
| STAR | 1.16% | 1.36% | 74149312 |
| LAST | 0.90% | 1.26% | 67015253 |
| Kallisto | 0.90% | 1.21% | 87667382 |
| HomeoRoq | 1.32% | 1.64% | 60946109 |
| EAGLE-RC | 0.54% | 0.80% | 57254486 |

Here, error is represented as a proportion of quantified reads rather than the proportion of mappable reads in the earlier analysis. Our results show that error decreased in STAR, LAST, and Kallisto as the quantification process accounted for ambiguously mapped reads in some fashion (dropped by featureCounts and distributed by Kallisto). HomeoRoq and EAGLE-RC error rate increased slightly due to these methods discarding reads that are deemed unclassifiable. However, a large number of unclassifiable reads are due to equivalent alignments to both subgenomes, deemed subgenome-common, which can be used to estimate expression levels such as RPKM by distributing proportionally to each subgenome. This has been demonstrated previously for HomeoRoq [30] and is also applicable to EAGLE-RC. Thus the number of reads quantified for subgenome-classification approaches in Table 2 may not be directly comparable to standard genome alignment approaches. It turns out that for tetraploid *A. kamchatica*, all methods had comparable error rates, where EAGLE-RC was the most precise.

Next, we examined the ratio of homeologous pairs. We are interested in any shifts in expression between homeologs across conditions, so we analyzed RNA-seq data from *A. kamchatica* and quantified the *A. halleri* read count proportion 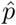 as follows:

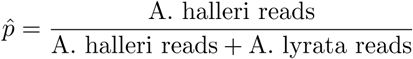

While we do not know the ground truth expression levels, we can evaluate the concordance between the different methods to describe how results might differ depending on which approach was used. To compare the results between different methods, we calculated the pairwise root mean squared distance (RMSD) and coefficient of determination (*r*^2^) between the results for all methods. The RMSD is a measure of the average distance between two sets *X* and *Y*:

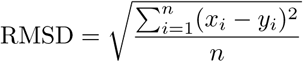

The *r*^2^ describes how well one variable *X* can be used to predict another variable *Y* by calculating the proportion of variability that can be explained:

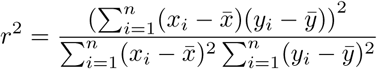

Though both measures describe the similarity between the results of different methods, the RMSD quantifies the magnitude of difference with units while the *r*^2^ quantifies the proportion of similar elements.

We examined the ratio of homeologous pairs using RNA-seq data from *A. kamchatica* samples before and after cold stress. The *r*^2^ (Table 3) and RMSD (Table S3) for 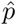 show that in general, the results from the standard genome alignment based approach were concordant (average *r*^2^ = 0.9505, RMSD of 6.96%). HomeoRoq was also concordant (average *r*^2^ = 0.9084) with STAR and LAST. This is consistent with the error rates in the classification results.

**Table 3:** *r*^2^ of the proportion of reads derived from the *A. halleri* subgenome between different quantification approaches for homeologs in *A. kamchatica*.

|  | STAR | LAST | Kallisto | HomeoRoq | EAGLE-RC |
| --- | --- | --- | --- | --- | --- |
| STAR | - |  |  |  |  |
| LAST | 0.9505 | - |  |  |  |
| Kallisto | 0.6886 | 0.6852 | - |  |  |
| HomeoRoq | 0.9214 | 0.8953 | 0.6472 | - |  |
| EAGLE-RC | 0.8201 | 0.8003 | 0.5791 | 0.8325 | - |

The generally high *r*^2^ between methods showed that though read mapping precision can vary greatly, downstream read counting methods were somewhat able to correct for artifacts or ambiguity in read alignment to arrive at a similar quantified expression. Kallisto was discordant from all other methods with an average RMSD of 18.88%. This may be due to its built-in read counting method using Expectation Maximization (EM) whereas all other methods used featureCounts, which does not consider multi-mapped reads. There were also issues with low expression genes where 420 and 447 *A. halleri* and *A. lyrata* homeologs respectively, were reported to have near zero expression (< 10 reads) by all other methods in all samples but Kallisto reported > 100 reads. The homeolog expression ratio scatter plots reflect this potential artifact (Figure S1). Though the cause is unclear, Kallisto tends to over-estimate some low expression genes compared to other methods.

EAGLE-RC also showed less concordance in *r*^2^ and RMSD than HomeoRoq to standard genome alignment approaches, where it reported some high expression genes as low expression compared to other methods. More detailed examination revealed that because EAGLE-RC required reads to cross genotype differences between homeologs, it excluded reads in exons which were not able to be pair-wise aligned. There may exist uncertainty in the annotation, as there are 552 homeologs that had at least a 40% difference in the proportion of the gene model aligned between *A. halleri* and *A. lyrata*. The majority of these cases were *A. halleri* homeologs having a smaller proportion than *A. lyrata* homeologs due to being much longer. The exclusion of regions which were not aligned between homeologs accounted for a large portion of the difference in EAGLE-RC. For example, the homeolog annotated as AT4G25110 in *A. kamchatica* with 12 exons in *A. halleri* and 5 exons in *A. lyrata* and an aligned region between homeologs of 46% and 61% in *A. halleri* and *A. lyrata* respectively, had almost all (99.89%) of the reads assigned to *A. halleri* mapped to regions that were not aligned between homeologs (Figure S2), leading to a large difference between EAGLE-RC and other methods (Table S4).

The subgenome-classification approach with HomeoRoq or EAGLE-RC tries to account for potentially missing reference genome regions in one subgenome, through consideration of read mappability and explicitly utilizing genotype variation respectively. For example, the homeolog annotated as AT5G45850 in *A. kamchatica* had reads that mapped to the *A. lyrata* reference that did not map at all to the *A. halleri* reference and was thus discarded by HomeoRoq. Similarly, missing reference genome regions in homeologs cannot be pair-wise aligned, thus EAGLE-RC discarded reads that map to these regions. In other methods, a naive counting of reads could be reference biased (Table S5). In this case, the average 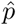 for a sample with the subgenome-classification approach was 0.90 compared to 0.44 with STAR, LAST, and Kallisto.

Next, we examined differentially expressed homeologs (DEH) using RNA-seq data from *A. kamchatica* samples before and after cold stress. The results of different methods (Figure 3) showed that in general, there was high overlap between the different methods tested where Kallisto was the least concordant among the methods tested. This is consistent with the results when we examined the ratio of homeologous pairs.

**Figure 3:**
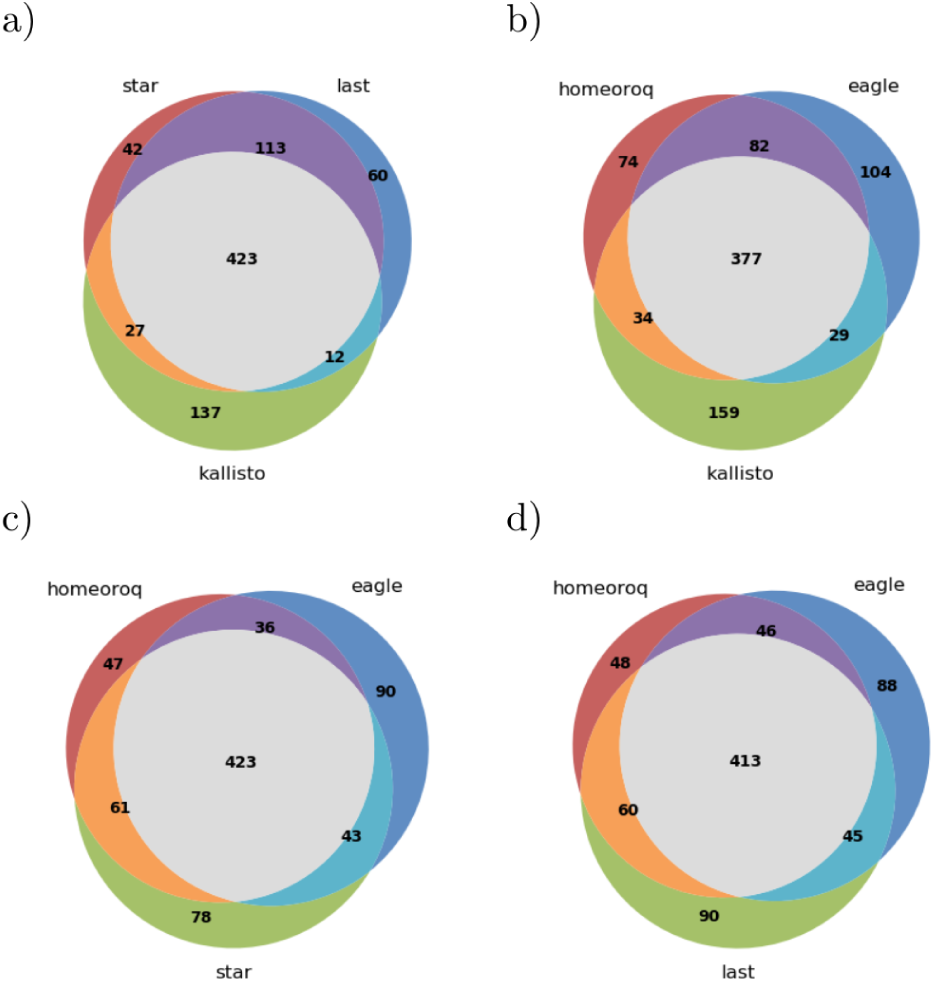
Overlap of differentially expressed homeologs from different methods in *A. kamchatica*.

Intuitively, genes with lower expression are expected to be more affected by uncertainties due to read count differences having more of an effect on the proportion (Table 4). Indeed, our results show that the concordance between methods was lower for genes with lower expression with a significant increase (paired t-test p-value 6.767 × 10 ^−06^) in RMSD. This may be due to a higher proportion of reads being ambiguous and each method’s difference in handling these ambiguously mapped reads. This may have large implications if the goal is to determine homeolog expression bias.

**Table 4:** RMSD of the proportion of reads derived from the *A. halleri* subgenome for homeologs with read counts ≤ 100 (lower triangular matrix) versus > 200 (upper triangular matrix) in *A. kamchatica*.

|  | STAR | LAST | Kallisto | HomeoRoq | EAGLE-RC |
| --- | --- | --- | --- | --- | --- |
| STAR | - | 0.0213 | 0.1008 | 0.0324 | 0.0457 |
| LAST | 0.1092 | - | 0.1008 | 0.0380 | 0.0499 |
| Kallisto | 0.2301 | 0.2301 | - | 0.1026 | 0.1076 |
| HomeoRoq | 0.1329 | 0.1529 | 0.2460 | - | 0.0380 |
| EAGLE-RC | 0.1897 | 0.2010 | 0.2725 | 0.1867 | - |

### 3.3 Hexaploid wheat read classification

For hexaploid wheat *T. aestivum*, the three homeolog copies compared to two in tetraploids further complicates read mapping and also requires a more complex workflow for subgenome-classification. To test classification performance, we used RNA-seq data of a tetraploid wheat line (tetra-Chinese Spring) with *A* and *B* subgenomes, and a diploid *A. tauschii* line (accession KU-2076) with *D* subgenome as the ground truth. Table 5 shows the classification performance of each approach.

**Table 5:** Classification performance for *T. aestivum*. Results are averaged across 3 samples for each of diploid *A. tauschii* (*D*) and tetraploid Chinese Spring (*AB*). The percent mapped refers to the number of reads that were mappable and in the case of subgenome-classification, the number of reads mappable to any subgenome. The classification error refers to the proportion of classified reads which were assigned to the wrong subgenome. The number of expressed homeologs with > 1 read in any sample are also shown for each subgenome.

|  |  | Classification error |  | chrA | chrB | chrD |
| --- | --- | --- | --- | --- | --- | --- |
|  | Mapped | AB to D | D to AB | Homeologs | Homeologs | Homeologs |
| STAR | 75.83% | 9.12% | 14.85% | 11366 | 11333 | 12530 |
| LAST | 77.21% | 1.62% | 2.78% | 11505 | 11437 | 12847 |
| Kallisto | 78.17% | 10.17% | 16.71% | 14794 | 14847 | 16032 |
| HomeoRoq | 79.73% | 0.97% | 1.54% | 10224 | 10217 | 11539 |
| EAGLE-RC | 79.73% | 0.33% | 0.66% | 10350 | 10285 | 11766 |

Our results show that the misclassification rate of all methods were strikingly similar for allotetraploid *A. kamchatica* and allohexaploid wheat (Figure 4). This suggests that it was mainly the difference in methods rather than taxon-specific features that affected the error rate where subgenome-classification methods showed a higher precision than standard methods using the concatenated genome. Higher precision was obtained in the order of EAGLE-RC, HomeoRoq, LAST, STAR, and Kallisto. For the number of expressed homeologs detected by these methods, Kallisto showed higher counts, which may be spurious because Kallisto tended to overestimate low expression genes relative to other methods as discussed above. Among the other four methods, the subgenome-classification methods showed a slightly lower number of expressed homeologs detected than the standard genome alignment approach, though we do not know which one is closer to the ground truth. EAGLE-RC still maintained a sub 1% error rate, the best among all methods, though it deemed an average of ~20% of mapped reads as unable to be classified with confidence due to ambiguity between homeolog pairs. There was also a higher cost in computation time for the higher precision (Table S6).

**Figure 4:**
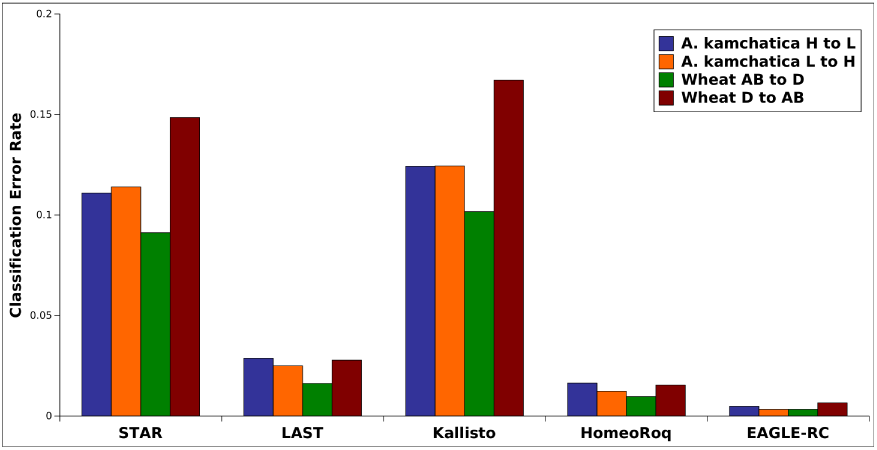
Overall classification error rate in tetraploid *A. kamchatica* and hexaploid wheat *T. aestivum* for all methods.

Another point of interest is that the more divergent *D* diploid line had much higher error rates than the *AB* line directly extracted from Chinese Spring, which is the reference genome line (Table 5). In this case, STAR and Kallisto performed quite poorly compared to other methods. Divergence from the reference genome appears to be a multiplier for error rate, thus less precise methods will be more affected.

### 3.4 Hexaploid wheat homeolog expression quantification

We evaluated the classification error using the quantified read counts for homeologs in *T. aestivum* (Table 6). As described in the previous section, note that the number of reads quantified are not directly comparable among different methods. Again, for Kallisto we used the estimated read count from its output while for all other methods we obtained read counts using featureCounts.

**Table 6:** Classification performance for *T. aestivum* using quantified read counts. Results are averaged across 3 samples for each of diploid *A. tauschii* (*D*) and tetraploid Chinese Spring (*AB*).

|  | Classification error |  | Reads quantified |
| --- | --- | --- | --- |
|  | AB to D | D to AB |  |
| STAR | 2.83% | 5.13% | 2632463 |
| LAST | 1.46% | 1.80% | 3324633 |
| Kallisto | 1.45% | 3.07% | 4922150 |
| HomeoRoq | 1.50% | 2.26% | 1929797 |
| EAGLE-RC | 0.67% | 1.32% | 1892448 |

As expected, the error rate was higher in hexaploid wheat than in *A. kamchatica* though with similar trends. In *AB* classification error, STAR had over ~200% higher rate of error, LAST and Kallisto had ~60% higher error, and HomeoRoq and EAGLE-RC were the most robust with ~20% higher error. Similar to the alignment error analysis, the more distant *A. tauschii* (*D*) reads had much higher error rates, propagated from errors in the alignment. Again, STAR and Kallisto were the most affected by the increased distance in these samples.

Next we examined homeolog expression shifts across conditions and quantified the homeolog expression ratio in *T. aestivum* by calculating the proportion of subgenome A reads 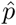 as follows:

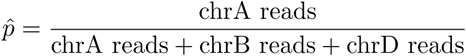

We examined the homeolog expression ratio using RNA-seq data from *T. aestivum* samples 24h after fungal inoculation and after mock inoculation. The *r*^2^ (Table 7) and RMSD (Table S7) show that relative to other methods, Kallisto exhibited a large drop in concordance in hexaploid wheat compared to *A. kamchatica*. The homeolog expression ratios (Figure 5) show that the discordance was largely due to systematic differences in low expression homeologs, which was also a trend for other methods (Figure S3-S5) though not to the degree of Kallisto. Also similar to *A. kamchatica*, there was some discordance between EAGLE-RC and other methods. In hexaploid wheat, there are 1689 *AB*, 1549 *BD*, and 1661 *AD* pair-wise homeologs that have at least a 40% difference in the proportion of the gene model aligned, which may account for much of the discordance between EAGLE-RC and other methods.

**Figure 5:**
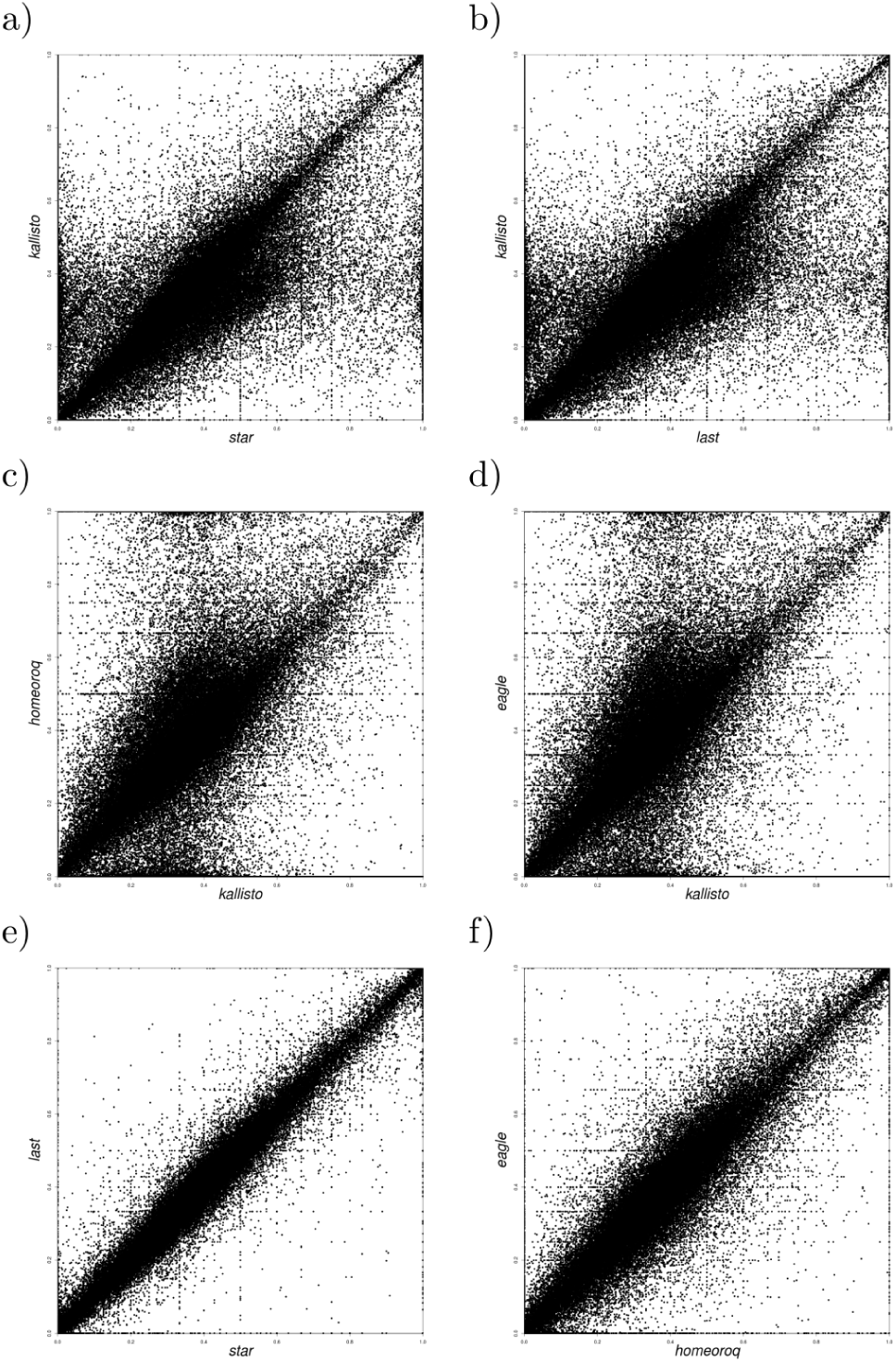
Homeolog expression scatter plots for Kallisto versus other methods in hexaploid wheat, quantified as the expression proportion of subgenome A over the total (*A*+*B*+*D*) per homeolog.

**Table 7:** *r*^2^ of the proportion of reads derived from subgenome *A* between different quantification approaches for homeologs in *T. aestivum*.

|  | STAR | LAST | Kallisto | HomeoRoq | EAGLE-RC |
| --- | --- | --- | --- | --- | --- |
| STAR | - |  |  |  |  |
| LAST | 0.9174 | - |  |  |  |
| Kallisto | 0.3257 | 0.3342 | - |  |  |
| HomeoRoq | 0.8612 | 0.8107 | 0.2787 | - |  |
| EAGLE-RC | 0.8310 | 0.7931 | 0.2814 | 0.7958 | - |

Next, we examined differentially expressed homeologs using RNA-seq data from *T. aestivum* samples 24h after fungal inoculation and after mock inoculation. The DEH results for hexaploid wheat between different methods (Figure 6) show that there was less concordance between methods compared to tetraploid *A. kamchatica*. Notably, only 51% of 304 DEH detected by Kallisto were supported by any of the other four methods (47% by STAR, 45% by LAST, 38% by HomeoRoq, 41% by EAGLE-RC). The results here reiterate that discordance was systematic, where Kallisto’s tendency to overestimate low expression reads led to significant differences in the homeolog expression ratio.

**Figure 6:**
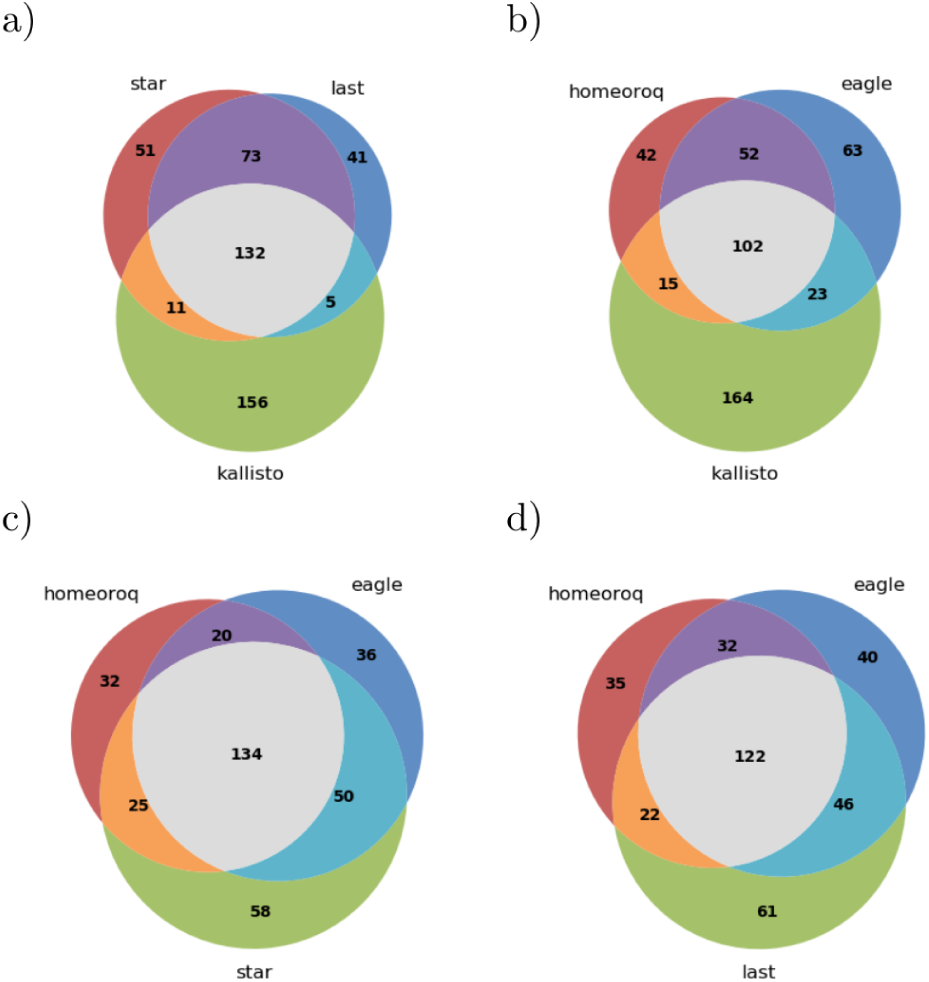
Overlap of differentially expressed homeologs from different methods in *T. aestivum*.

## 4 Discussion

Recent improvements in sequencing technology has reduced the difficulty in constructing allopolyploid reference genomes [12, 13, 14] and there should be a corresponding increase in genome wide studies for allopolyploids [3, 4, 5, 6, 7, 8]. As such, there is a need to evaluate expression quantification methods given the presence of homeologs in allopolyploids. Unlike expression quantification in diploids, homeolog expression quantification evaluates multiple highly similar gene copies concurrently and it would be ideal if all copies are at a similar level of completeness. This includes the genome annotation, which may be a non-trivial source of uncertainty [36, 37].

It is well known that the presence of repetitive sequences in diploids (paralogs) present technical challenges for read alignment [38, 39] and can bias RNA-seq expression quantification [40]. For polyploids, with the presence of homeologs, there are even more repetitive sequences due to an increase in the number of gene copies. In this study, we saw that applying a standard diploid RNA-seq workflow to allopolyploids may have issues in terms of assigning reads to the wrong subgenome, particularly in hexaploid wheat. In general, low expression genes accounted for most of the discordance between methods. Kallisto especially, often overestimated the number of reads in low expression genes which has been observed in previous studies [41]. If the goal is to determine homeolog expression bias, then accurate quantification of low expression genes is important because small errors can result in large shifts in the expression ratio between homeologs. In addition, the accuracy of expression levels in such genes is especially important if we wish to identify ON and OFF states of gene expression in response to stimuli in time course analyses.

There has long been a discussion and dispute about the bias of expression between homeologs and subgenomes because different species show different patterns [42]. In this study we found that different methods showed different biases, depending on the types of uncertainty they consider. In particular, gene annotation can affect the detection of homeolog expression bias because the exon regions of homeologs are typically annotated using RNA-seq reads on each copy separately and thus may be annotated differently, especially when the expression level of one of the copies is low. To obtain general conclusions on the biased expression in polyploids, we would suggest that analysis be performed with comparable and accurate methods in corresponding exonic regions. EAGLE-RC attempts to do this by considering only corresponding exonic regions between homeologs.

In our evaluations, we presented results both at the read mapping stage and after expression quantification. It is difficult to be completely fair when evaluating the accuracy of read mapping due to differences in each of the four approaches. Kallisto does not attempt to identify primary versus secondary alignments, thus a primary alignment only evaluation is not suitable. The standard genome alignment approach may have secondary alignments due to the presence of homeologs, which can be indistinguishable from the primary alignment in terms of alignment score, in which case a primary alignment is picked randomly. The subgenome-classification approach of mapping reads to each subgenome separately inherently has less secondary alignments, which is an advantage of this approach. Thus we evaluated the classification accuracy with all alignments, as secondary alignment errors are a factor in all approaches though to different degrees. The results after read counting then presents the performance of each approach after potentially accounting for ambiguous alignments. In this way we show the performance at both major steps in the RNA-seq expression quantification process, though there are many other read counting methods that we were not able test. Read counting methods may also benefit from the higher precision of the subgenome-classification approach by utilizing classified reads in lieu of unique mapped reads and distributing ambiguous reads through Expectation Maximization.

## 5 Conclusion

In this study, we evaluated methods for homeolog expression quantification in tetraploid *A. kamchatica* and hexaploid wheat *T. aestivum* using RNA-seq. We examined the standard genome alignment based approach with STAR and LAST, the subgenome-classification approach with HomeoRoq and EAGLE-RC, and a pseudo-alignment approach with Kallisto.

The presence of homeologs had the largest affect on STAR and Kallisto, resulting in higher read classification error. We observed that discordance occurred mostly in low expression genes in a systematic manner and can result in large shifts in the homeolog expression ratio. The explicit use of genotype differences between homeologs in EAGLE-RC seems to be a factor in reducing uncertainties in the reference genome and annotation and our results show that EAGLE-RC was the most precise method in both tetraploid *A. kamchatica* and hexaploid wheat.

## Author Contributions

T.K. executed the study and drafted the manuscript. M.H. contributed to data analysis and simulation studies. T.T. contributed to genetic material and data collection. K.K.S and J.S. initated the study, the analysis framework, and contributed to drafting the manuscript.

## Acknowledgments

The *Aegilops tauschii* seeds were kindly supplied by the National BioResource Project-Wheat (Japan, http://www.nbrp.jp). The tetra-Chinese Spring seeds were kindly supplied by Professor Hisashi Tsujimoto (Tottori Univ.). Dr. Yoshihiro Matsuoka (Fukui Prefectural Univ., Japan), Dr. Hiroyuki Tsuji (Yokohama City Univ.) and Ms. Akina Mitsuhashi (Yokohama City Univ.) provided generous support for *Aegilops tauschii* cultivation and propagation. We thank Rie Shimizu-Inatsugi for plant experiments, Catharine Aquino and the Functional Genomics Center Zurich for sequencing, Thomas Wicker, Beat Keller, and Cristobal Uauy for valuable discussions.

